# A Transcriptional Atlas of Endothelial Cell Zonation Along the Pulmonary Vascular Tree

**DOI:** 10.1101/2025.05.17.654540

**Authors:** Stefanie N. Sveiven, Carsten Knutsen, Fabio Zanini, David N. Cornfield, Cristina M. Alvira

**Affiliations:** Division of Critical Care Medicine, Department of Pediatrics, University of California San Francisco, San Francisco, CA 94158, USA; School of Clinical Medicine, UNSW Sydney, 2052, NSW, Australia; UNSW Cellular Genomics Futures Institute, 2052, NSW, Australia; UNSW Evolution & Ecology Research Centre, 2052, NSW, Australia; Center for Excellence in Pulmonary Biology, Stanford University School of Medicine, Stanford, CA 94305, USA; Division of Pulmonary, Asthma and Sleep Medicine, Department of Pediatrics, Stanford University School of Medicine, Stanford, CA 94305, USA

## Abstract

**Background:** The lung vasculature is comprised of a series of branching vessels extending from the main pulmonary artery to the alveolar capillaries, then back to the pulmonary veins. Lung endothelial cells (EC) exist along this continuum, exposed to gradients of shear stress, oxygen tension and pressure. Single cell RNA sequencing (scRNA-seq) has identified lung EC subsets, but many aspects of the vascular continuum, including vessel size and capillary polarity remain undefined from transcriptomic data.

**Methods:** We created an endothelial-enriched scRNA-seq dataset from the P3 mouse lung. Using diffusion pseudotime across all lung EC, we developed an analytical framework to delineate transcriptomic gradients and assign vessel-size scores to categorize individual endothelial cells (EC) along the vascular continuum. We confirmed size-related gene expression patterns with fluorescence in situ hybridization.

**Results:** We categorized capillary 1, arterial and venous EC along two gradients: arterio-venous zonation and vessel size. This approach distinguished large arteries from arterioles, large veins from venules, and revealed arterio-venous polarity within the capillaries. Our data recapitulated previously established zonally defined cell signaling axes, such as high *Cxcl12*-*Cxcr4* signaling in arterioles. We also identified unique cellular communication occurring in large versus small arteries and veins, and localized injury-induced venous EC proliferation to vessels of specific size. This analytical framework was successfully applied to several published mouse and human datasets across different stages of lung development.

**Conclusions:** These findings provide a comprehensive transcriptional map of EC across the pulmonary vascular tree, enabling assignment of each individual cell to vessels with defined size and position. This framework offers spatial inferences and novel mechanistic insights from scRNA-seq data sets that may elucidate therapeutic targets to treat pulmonary vascular diseases affecting specific vascular segments. We speculate that similar frameworks could be applied to tissues outside the lung.

## Introduction

Recent advances to profile the transcriptomes of single cells have transformed our understanding of biology, revealing previously underappreciated diversity across cellular subtypes, including the heterogeneity of pulmonary vascular cells during development and in response to injury^1–6^. Although most researchers categorize cell states into discrete clusters^7^, cell states often exist along continuums. Breaking these continuums into arbitrary groups may conceal important biological signals. For example, pulmonary endothelial cells (EC) lining the vascular continuum receive distinct physiologic signals (e.g. blood flow, pressure, pulsatility, and oxygen tension) depending on their precise location along the arterial-capillary-venous axis^8^.

Thus, broad categorization of EC as artery, vein, or capillary likely obscures transcriptomic and phenotypic variation of cells across these physiological axes^9,10^.

The entire cardiac output passes through the lungs with each heartbeat, allowing all the deoxygenated blood to pass through gas exchanging alveoli to ensure distribution of oxygen rich blood to the remaining organs. Pulmonary arterial (PAEC), capillary and venous (PVEC) EC lining the pulmonary circulation play distinct roles: modulating vascular tone to optimize ventilation and perfusion, performing gas exchange while maintaining a tight barrier to prevent fluid extravasation into the alveolus, and regulating leukocyte trafficking and immune responses to pathogens entering through the alveolar space^11^. In addition to their primary role in transporting oxygen and nutrients, EC also supply angiocrine signals that inform local cellular niches and influence the behavior of neighboring cells, regulating tissue development and repair^8^.

Although single cell transcriptomics has identified distinct gene expression signatures exhibited by broad pulmonary EC subtypes^9,10^, EC located at proximal versus distal positions of the circulation encounter distinct microenvironments. Given that many pulmonary vascular diseases selectively affect specific locations within the circulation, developing computational methodologies to identify transcriptomic changes in pulmonary EC at specific locations will enhance our ability to understand alterations during lung development and disease^12,13^. Moreover, computational methods to infer cell-cell communications from single-cell data often propose interactions between cells that are spatially distant and therefore unlikely to directly interact.

Transcriptional information on endothelial cell zonation can provide spatial information that increases the probability that specific cells are physically capable of communication.

Using a strategy of endothelial enrichment and single cell RNA sequencing to create a high resolution, transcriptomic dataset from the developing mouse lung, we developed an analytical framework to assign vessel-size scores and categorize individual EC along a continuum of vessel size. We delineated a continuum of arterial to venous, and macro-to microvascular zones using these sized-based transcriptional signatures. Our strategy allowed identification of signaling axes previously established with spatial methodologies solely from transcriptomic data, and localization of disease relevant alterations in gene expression to specific segments of the vasculature. This vessel-size informed framework is robust across development and species and reveals how spatial EC heterogeneity underlies key processes in lung development and injury.

## Materials and Methods

### Sample preparation

At birth, neonatal C57BL-6 (Charles River) mice were housed in room air (normoxia; N) or 80% O_2_ (hyperoxia; H) for 72hr before euthanasia (N=6 per exposure; 3M 3F). Lungs were perfused with HBSS, excised, finely minced, pooled, and digested in Liberase TM (0.2mg/mL) plus DNaseI (0.01mg/mL) for 10min at 37.5 in a bacteriological shaker. The suspension was triturated 15 times. Liberase was inactivated using 2mM EDTA and cold FBS prior to pelleting the cell suspension. Cells were treated with 1X RBC Lysis for 5min then passed over a 30μm cell strainer before microbead enrichment (SmartStrainer, Miltenyi). In a parallel set of experiments, lungs were pressure fixed at 20cm H_2_O and paraffin embedded (FFPE) for *in situ* validation experiments.

### Enrichment by magnetic associated cell sorting (MACS)

Immune cells were depleted using anti-CD45 coated Dynabeads (Invitrogen). The supernatant was pelleted, counted, and resuspended with blocking buffer (anti-mouse CD16/CD32 and anti-rat IgG; 1:100) for 10min. Mouse anti-BST1–APC (PVEC marker; Biolegend, clone BP-3) and rat anti-CD31–biotin (pan-EC marker; BD Pharminogen, clone MEC 13.1) antibodies were added to the cells (2μg/10M cells) and incubated on a HulaMixer for 20 minutes. To ensure sufficient PVEC for our analysis, following the manufacturer’s protocol, cells were first enriched for BST1 using the MACS anti-APC microbeads and then the flow through was enriched for CD31 using anti-biotin microbeads (Miltenyi).

### Chromium GEMX 3’ v4 gene expression

Following the manufacturers protocol, enriched populations were then adjusted to 1500 cell/μL to obtain the 20,000 target cell input for gel bead emulsion formation (10X Genomics). Samples that met QC standards (Bioanalyzer, Stanford Protein and Nucleic Acid core) were used for barcoding, library construction and sequenced by NovoSeq Χ at a depth of 1B paired reads per sample (NovoGene Corporation Inc.).

### scRNA-seq Analysis

Sequencing reads were aligned to the Grcm39 mouse genome using Cellranger (v9.0.0). SoupX was used to remove ambient RNA signal^14^. Gene expression count tables were turned into anndata objects and processed using scanpy when not otherwise specified^15^. Cells that had more than five median absolute deviations of either unique genes or UMIs detected were removed. Cells that had more than three median absolute deviations of percent mitochondrial or ribosomal UMIs were removed^7^. Scrublet was used to automatically detect and remove doublets. Counts were normalized and log transformed^16^. PCA was run prior to embedding and clustering. The Leiden algorithm was used for clustering, and Uniform Manifold Approximation and Projection for embedding. Lineage identity was assigned using canonical markers: *Cdh5* (endothelial); *Epcam* (epithelial); *Ptprc* (immune); and *Col1a1* (mesenchymal). Cell typing was performed after each lineage was re-clustered and embedded as detailed above. Cell types were assigned a second time after regressing out cell cycle genes^7^. Pseudotime analysis and branching was performed using Palantir^17^, and RNA velocity determined using velocyto to align reads, and scVelo to calculate trajectories^18,19^. Vessel size scoring was performed by finding shared genes within the top 50 genes positively or negatively correlating, using Pearson’s correlation, with pseudotime in PAEC or PVEC, using Cap1 EC as the root. Scores were generated for positively and negatively correlated genes separately, and the negative correlation score subtracted from the positive score, then scaled to be between 0-1. Vessel size categories were assigned from quartiles in the vessel size score. For external datasets, Cap1, PAEC and PVEC were subset from the dataset using the original annotation. These cells were re-embedded and clustered to assign consistent cell type names across datasets and remove low-quality cells. Vessel size scoring was done the same as described above. For more detail see https://github.com/CarstenKnutsen/Vessel_size_manuscript

### Validation by RNA *in situ* hybridization and immunofluorescence

RNAScope MultiPlex v2 Assay (Advanced Cellular Diagnostics) and immunofluorescence was performed on 5μm FFPE lung sections from P3 mice. RNAScope probes were purchased from ACD and fluorophores from Akoya: Neonatal mice were given a single dose 20mg/kg of intragastric EdU dissolved in PBS 2hours before tissue collection. ClickiT EdU was performed after completion of the RNAScope protocol per the manufacturer’s instructions. Fluorescence images were captured using a Zeiss Axio Observer 7 equipped with Apotome for optical sectioning using a 20X objective. Images were quantified using CellProfiler (v4.2.6)^20^. A minimum of 10 images were taken per mouse, n=3-5 per group. From these images, vessels were cropped using the “IdentifyObjectsManually” module. Each cropped vessel was then measured for their signal intensity from each channel was using the “MeasureImageIntensity” module. For EdU+ PVEC quantification, nuclei, EdU signal and Slc6a2 signal were identified using the “IdentifyPrimaryObjects” module. Overlap of these signals was done using the “RelateObjects” module. Vessel diameter was determined using the outer diameter of the major axis.

### Quantification and statistical analysis

To identify differentially expressed genes Wilcoxon rank sum tests were run on all genes’ expression between cell populations, with a false discovery rate (FDR) adjustment with Benjamini-Hochberg, significance was determined as having a FDR<0.05. Differences between two groups in image analysis were determined by Student’s t-test, and correlations were done using a Pearson correlation.

## Data Availability

Data generated for this manuscript is deposited at the Gene Expression Omnibus under GSE315745. Postnatal mouse data was sourced from the Gene Expression Omnibus under code GSE151974. Adult mouse data was sourced from https://datasets.cellxgene.cziscience.com/e818d27c-62ae-4c19-97c7-6cd4d65b8f9b.h5ad. Human neonatal data was sourced from https://cellxgene.cziscience.com/collections/28e9d721-6816-48a2-8d0b-43bf0b0c0ebc. Human 1 month-3-year-old data was sourced from https://www.lungmap.net/dataset/?experiment_id=LMEX0000004400. Public adult human data was sourced from https://figshare.com/articles/dataset/Tabula_Sapiens_v2/27921984.

## Results

### Pulmonary endothelial cells exhibit gradients of gene expression that span macro-and microvascular transitions and delineate pre-and post-alveolar capillaries

Pulmonary vascular EC exist along a structural continuum of large to small arteries and veins, including macrovascular to microvascular transitional zones where general capillaries/capillary 1 (gCap/Cap1) flank gas-exchanging aerocyte capillaries (aCap/Cap2) on either the pre-alveolar (arterial) or post-alveolar (venous) side of the circulation (**Fig. 1A**). To delineate EC diversity among the pulmonary vasculature, we performed single cell RNA sequencing (scRNA-seq) on pulmonary cells from mice at the saccular stage of development, a period of rapid vascular growth. Endothelial cells, from mice exposed to either normoxia or hyperoxia, an injury that disrupts microvascular growth and induces pathologic vascular remodeling^3,21^, were enriched to ensure representation of small subpopulations^3,21^. A total of 32,946 cells representing all lineages and cell types (**Fig. 1B; SFig. 1A)**, including 11,724 EC, were profiled with a median of 4,586 genes detected per cell. In the global lung UMAP (**Fig. 1C**), Cap1 embedded between PAEC and PVEC consistent with the physical proximity to macrovessels at pre-and post-capillary macro-micro transition zones^1,6^. Re-embedding these three populations provided additional resolution of these macro-microvascular transitions with a central Cap1 cluster positioned between PAEC and PVEC populations (**Fig. 1D)**, paralleling the structure of the pulmonary vascular tree.

**Figure 1:**
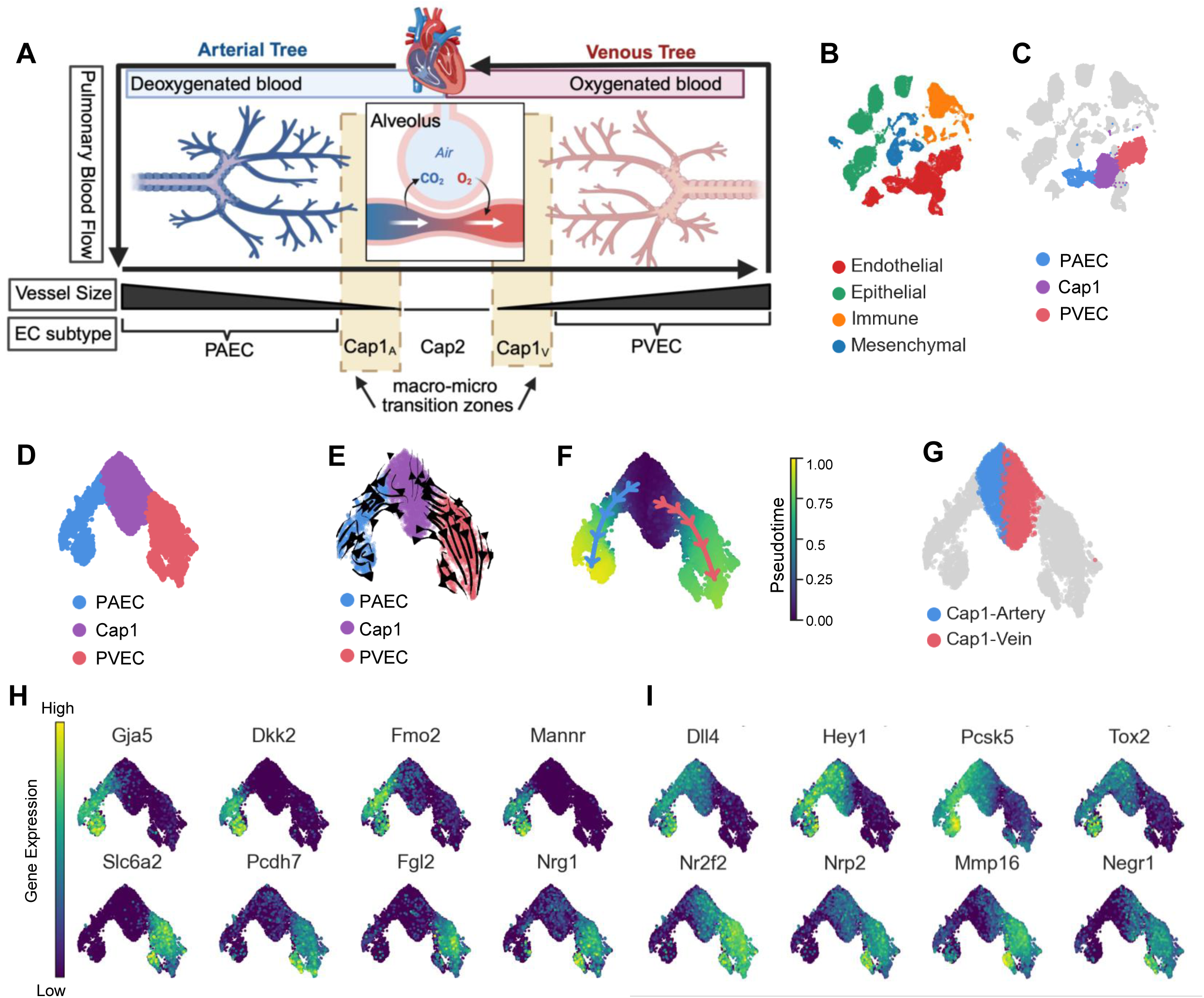
Pulmonary endothelial cells exhibit gradients of gene expression that span macro-and microvascular transitions and delineate pre and post alveolar capillaries. (A) Diagram of the lung vasculature demonstrating gradients of vessel size extending from the right heart through the pulmonary arteries, through progressively smaller vessels until gas exchange at the alveoli and venous size increasing throughout with return to the heart [made with BioRender]. (B) UMAP of all cells showing assigned lineage contingent upon expression of either *Cdh5* (endothelial); *Epcam* (epithelial); *Ptprc* (immune); or *Col1a1* (mesenchymal). (C) UMAP of vascular EC cell types highlighted: PAEC (blue), Cap1 (purple), and PVEC (red). (D) UMAP of PAEC, Cap1 and PVEC embedded alone. (E) UMAP colored by vascular EC cell type with RNA velocity vectors overlayed. Arrow of vector points in the direction of cellular trajectory. (F) UMAP colored by pseudotime (low to high: purple to yellow) with predicted branches overlayed. (G) UMAP with only Cap1 colored based on their arterio-venous sidedness. (H) UMAP feature plots of expression of arterial and venous marker genes (low to high: purple to yellow) showing markers largely restricted to macrovasculature, or (I) extending into polarized Cap1 EC.

To investigate the dynamic progression of EC states along the vascular continuum, we performed trajectory inference analysis. RNA velocity vectors inferred that Cap1 cells can progress towards either PAEC or PVEC (**Fig. 1E**), while branched pseudotime analysis revealed progressively increasing pseudotime values from the Cap1 node towards the far edges of either PAEC or PVEC nodes **(Fig. 1F)**. This pattern suggested that spatial localization of Cap1 cells as either pre-alveolar (Cap1_A_; arterial-adjacent) or post-alveolar (Cap1_V_; venous-adjacent) could be inferred from the transcriptome **(Fig. 1G).** When examining the expression of arterial and venous marker genes, some canonical arterial (*Dkk2*, *Gja5*) and venous markers (*Slc6a2*) were confined to their respective clusters, exhibiting little to no expression among Cap1 (**Fig. 1H**). However, other arterial and venous markers were enriched within each half of the Cap1 cluster, including elevated expression of canonical arterial (*Dll4*, and *Hey1*) or venous markers (*Nr2f2*, and *Ackr3*) within these putative Cap subsets **(Fig. 1I)**. Altogether, these data highlight transcriptional gradients of arterio-venous zonation, unique gene expression patterns among the vascular hierarchy, and a polarization of transcriptomic identities among Cap1 identified from our high-resolution scRNAseq dataset.

### The transcriptomic continuum of pulmonary endothelial cells aligns with vessel size

The transcriptional polarization of Cap1 towards PAEC and PVEC nodes suggested that an endothelial transcriptional continuum may parallel the structural branching of the lung vasculature. One characteristic of pulmonary vascular patterning is the gradual decrease in the diameter of vessels as proximal vessels successively branch distally toward the capillaries, with associated differences in matrix and mural cell coverage, providing additional transcriptomic clues to infer vessel size. Thus, we next identified genes negatively and positively correlated with pseudotime for both the PVEC and PAEC, with approximately one-third of these genes shared between both groups (**Supplemental Fig. 2A**). Notable examples of genes correlated with increasing pseudotime included elastin (*Eln*) and von Willebrand Factor (*Vwf*) known to be expressed by macrovessels. In contrast, microvascular markers like proto-oncogene (*Kit*) and transmembrane protein 100 (*Tmem100*), were among the genes that negatively correlated with pseudotime **(Fig. 2A)**^6,22^.

**Figure 2:**
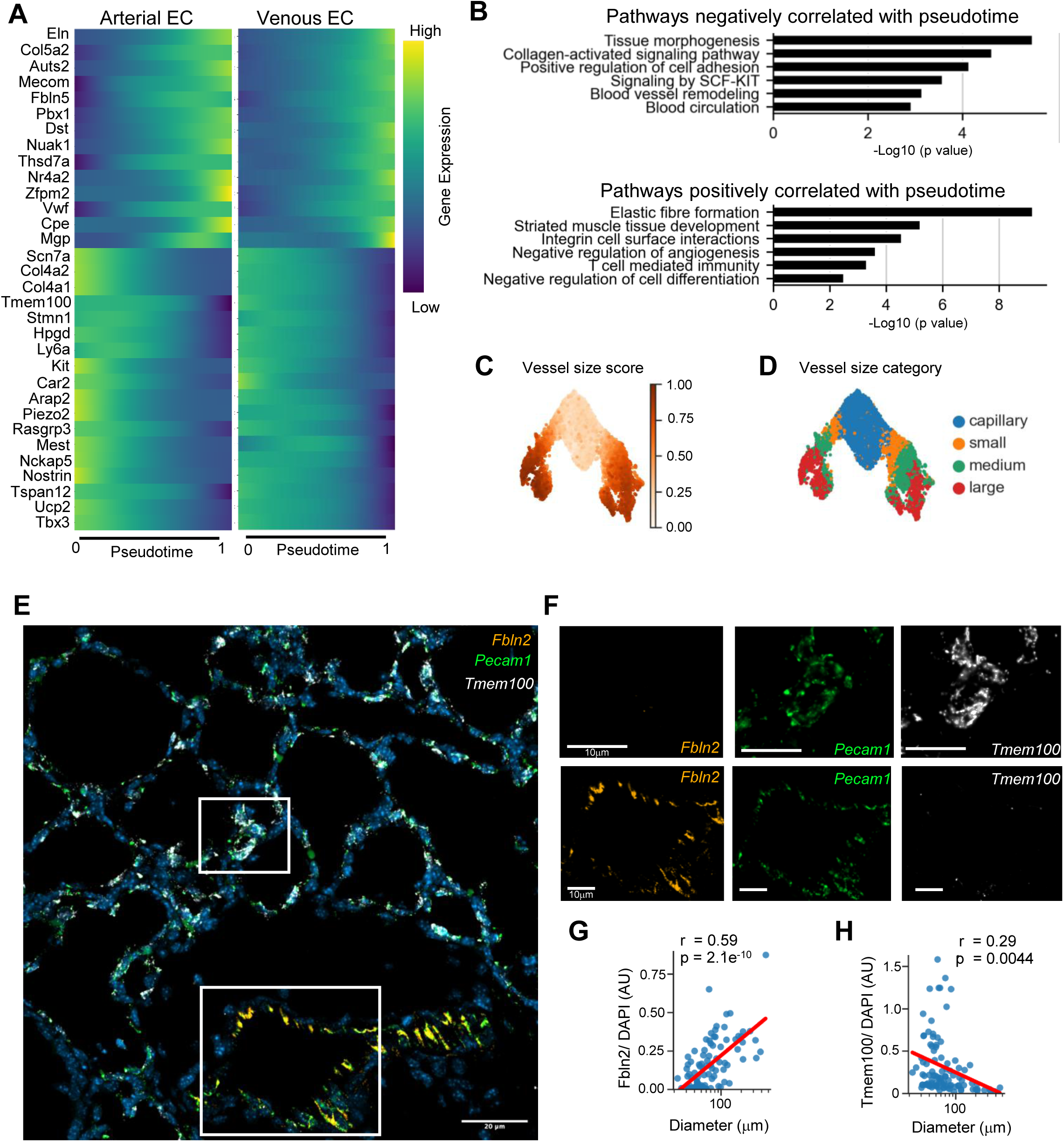
**The transcriptomic continuum of pulmonary endothelial cells aligns with vessel size size can be inferred in ECs from pulmonary vasculature**. (A) Heatmaps of top, shared genes correlated with pseudotime in both arterial and venous branches. Pseudotime increases from left to right in each heatmap, with low gene expression in purple and high gene expression in yellow. (B) Pathway analyses showing select enriched pathways with shared pseudotime correlation. (C) UMAP of vessel size score based upon expression of genes in panel A, with low size score in white and high size score in orange. (D) UMAP colored by vessel size category, with size quartiles determined by the vessel size score. (E) Representative image of multiplexed *in situ* hybridization to detect expression of the endothelial marker gene *Pecam1* (green) large vessel marker *Fbln2* (red) and small vessel gene *Tmem100* (yellow) in lung tissue from mice at P3 in normoxia. Calibration bar=20 μm (F) High magnification images of selected small (top) and large (bottom) vessels, showing each gene in separate images. Calibration bar=10 μm. Blinded quantification of (G) *Fbln2* and (H) *Tmem100* signal normalized to DAPI signal in vessels of various vessel sizes. Line in red is a linear regression on the log10 normalized vessel size, r and p value are from Pearson’s correlation. Each point represents a vessel. Vessels were imaged from 3 mice, with a total of n=97 vessels.

Pathway analysis on the top positively and negatively correlated genes with pseudotime also supported size-dependent physiology and function **(Fig. 2B; SFig. 2B)**. Developmental and angiogenic pathways such as ‘tissue morphogenesis’, ‘blood vessel remodeling’, and ‘signaling by SCF-KIT’ were enriched in negatively correlated genes, while positively correlated genes included enrichment of pathways include ‘elastic fiber formation’, ‘striated muscle development’, and ‘negative regulation of angiogenesis’-pathways aligned with the structural and functional characteristics of larger vessels.

We next used these shared pseudotime-correlated genes to calculate vessel size scores for each individual EC (**Fig. 2C**). Projecting these vessel-size scores onto the UMAP revealed a progressive distribution of vessel sizes along the vascular hierarchy, with the largest scores embedding along the terminal ends of the macrovascular clusters and the lowest scores embedding on the Cap1 node. We sorted scores into four bins representing ‘large’, ‘medium’, ‘small’, and ‘capillary’ based on quartile score ranges (**Fig. 2D**). The size nomenclature enabled us to visualize the assigned size distributions along the UMAP, such that the ‘capillary’ size embedded fully on the Cap1 cluster; the ‘small’ bin comprised the arms of PAEC and PVEC nodes connecting to the Cap1 cluster, while the ‘medium’ and ‘large’ cluster extended progressively outwards toward the distal edges of each macrovessel node.

We then validated this transcriptomic scoring of vessel size using RNA fluorescence *in situ* hybridization. We detected the expression of a small vessel marker, *Tmem100*, and a large vessel marker, *Fbln2,* in vessels of varying diameters (**Fig. 2E**). Images demonstrated the high expression of *Fbln2* but minimal *Tmem100* in larger vessels, and the reciprocal pattern in small vessels. Quantification confirmed that *Fbln2* expression was positively correlated with vessel size, with increasing signal as vessel diameter increased **(Fig. 2F)**. Conversely, *Tmem100* exhibited a negative correlation with diameter **(Fig. 2G)**. Together, these data support that our vessel-size analytical framework enables us to infer the relative size of the vessel from which an EC is derived based on gene expression, providing insights into spatial organization solely from transcriptomic data.

### Marked size-dependent transcriptomic heterogeneity within endothelial subtypes

We next identified genes exhibiting size-related expression patterns common to both arteries and veins (**Fig. 3A**). EC from the largest arteries and veins exhibited high expression of *Nr4a2*, an orphan nuclear receptor involved in vascular homeostasis and maturation, matrix Gla protein (*Mgp)*, an essential inhibitor of vascular calcification and elastin (*Eln*), a protein providing elasticity to larger vessels^23–25^. Medium sized PAEC and PVEC highly expressed the endothelial quiescence transcription factor *Foxo1*; G-coupled protein receptor *Adgrg6,* which promotes angiogenesis by modulating VEGF signaling; and receptor protein tyrosine phosphatases *Ptprr* ^26–28^. Small vessels had the highest expression of interferon-induced antiviral, transmembrane protein *Ifitm3*^29^; neuropilin-1 (*Nrp1*) a transducer of VEGF signaling that mediates angiogenic behavior^30^; and the transcription factor *Sox4*. Lastly, Cap1 EC highly express glucagon-like peptide-1 receptor (*Glp1r)*, *Ccdc85a*, a gene which supports barrier function, and *Kit*, a gene expressed by progenitor cells and a known Cap1 marker^1,3,6^.

**Figure 3.**
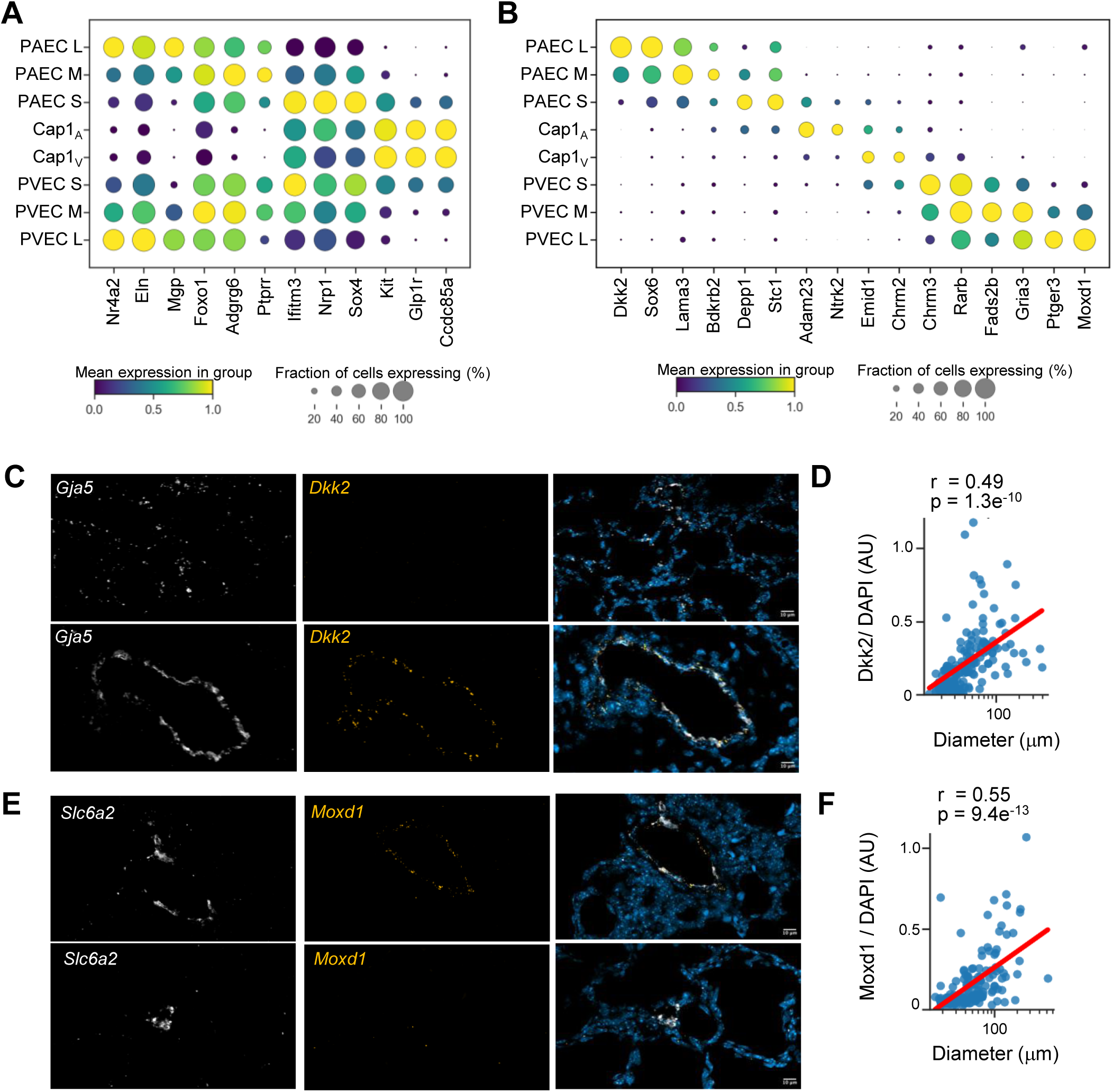
Marked size-dependent transcriptomic heterogeneity within endothelial subtypes. (A) Dotplot showing gene expression associated with size shared across arteries and veins, with relative gene expression indicated by color (low to high: purple to yellow), and percent of cell population expressing each gene indicated by the size of the circle. (B) Dotplot showing gene expression of gene expression associated with size that are distinct among arteries and veins, with relative gene expression indicated by color (low to high: purple to yellow), and percent of cell population expressing each gene indicated by the size of the circle. (C) Representative image of multiplexed in situ hybridization to detect expression of the arterial marker gene *Gja5* (white) and the large artery marker *Dkk2* (yellow) in lung tissue from mice at P3 in normoxia. Calibration bar=10 μm. (D) Quantification of *Dkk2* signal normalized by DAPI signal by vessel size. Each point represents a vessel. Vessels were imaged from 5 mice, with a total of 152 vessels. Line in red is a linear regression on the log10 normalized vessel size, r and p value are from Pearson’s correlation. (E) Representative image of multiplexed in situ hybridization to detect expression of the venous marker gene *Slc6a2* (white) and the large vein marker *Moxd1* (yellow) in lung tissue from mice at P3 in normoxia. (F) Quantification of *Moxd1* signal normalized by DAPI signal by vessel size. Each point represents a vessel. Vessels were imaged from 4 mice, with a total of 144 vessels. Line in red is a linear regression on the log10 normalized vessel size, r and p value are from Pearson’s correlation.

We then identified genes with distinct size-related expression patterns across the arterial and venous sides of the pulmonary circulation **(Fig. 3B)**. Consistent with our previous report, PAEC from large arteries showed highest expression of the Wnt-inhibitor *Dkk2*^3^ and the transcription factor *Sox6.* PAEC from medium sized vessels were enriched for *Lama3*, a laminin involved in basement membrane organization^31^, and *Bdkrb2,* encoding the bradykinin receptor B2, a regulator of vascular tone^32^. PAEC from small arteries were enriched for the HIF-responsive remodeling and stress-response genes *Depp1* and *Stc1*, Cap1 on the arterial side were enriched for *Adam23,* a regulator of integrin-mediated adhesion^33^, and *Ntrk2* a gene uniquely induced in adult Cap1 following injury^34^. Cap1 on the venous side expressed *Emid1*^35^, and the muscarinic receptor, *Chrm2* a regulator of vasomotor tone and angiogenic potential^36^. Among PVEC derived from small vessels, the muscarinic cholinergic receptor 3 (*Chrm3*,) was increased^37^. EC from both small and medium sized veins highly expressed the retinoic acid receptor *Rarb*, a component of a pathway known to promote lung vascular and alveolar development^38,39^. *Fads2b*, a fatty acid desaturase important for polyunsaturated fatty acid metabolism, and glutamate receptor *Gria3*, were also up-regulated in EC from medium veins. PVEC ascribed to large veins were enriched for the tumor suppressor *Moxd1*, and *Ptger3,* encoding the prostaglandin E receptor 3(**Fig. 3B)**.

We validated these size specific gene expression signatures in arteries and veins using *in situ* hybridization. In arteries, we detected the large arterial marker, *Dkk2*, in combination with the common arterial EC marker gene, *Gja5* (**Fig. 3C**). Imaging confirmed high expression of *Dkk2* restricted to large arteries, and quantification confirmed these results, demonstrating a positive correlation of *Dkk2* with increasing vessel diameter **(Fig. 3D)**. Using the same approach in veins to identify the common venous EC marker, *Slc6a2* with the larger vein marker*, Moxd1,* we found high expression of *Moxd1* restricted to large veins, with quantification showing a strong positive correlation with increasing venous diameter (**Fig 3E-F)**. Taken together, these findings demonstrate that even within canonical lung endothelial subtypes, there is marked regional transcriptomic heterogeneity of EC corresponding to specific locations along the vascular tree.

### Transcriptomic determination of vessel size provides biologic insight into endothelial-derived cellular communication

We next further validated our vessel-size analytical framework by determining if it could identify alterations in molecular signaling affecting specific segments of the pulmonary circulation previously identified using histologic data. For example, autocrine CXCL12-CXCR4 signaling promotes arteriolar patterning and branching during lung development^40,41^. Consistent with these data, we found the highest expression of *Cxcl12* and *Cxcr4* expression in small and medium sized PAEC (**Fig. 4A**). High *Ackr3* expression, a decoy receptor that clears excess CXCL12, was found in PVEC from large veins, suggesting a role for pulmonary veins in restricting CXCL12-CXCR4 signaling to the pulmonary circulation (**Fig. 4B**). Dysregulated CXCL12-signaling is implicated in the pathobiology of PH, a disease which entails pathologic remodeling primarily affecting small, resistance pulmonary arteries. Recent evidence suggests that estrogen may regulate *Cxcl12* expression^41–44^ and abrogation of Esr2 reduces muscularization of small-medium arteries in experimental models of PH^45^.

**Figure 4.**
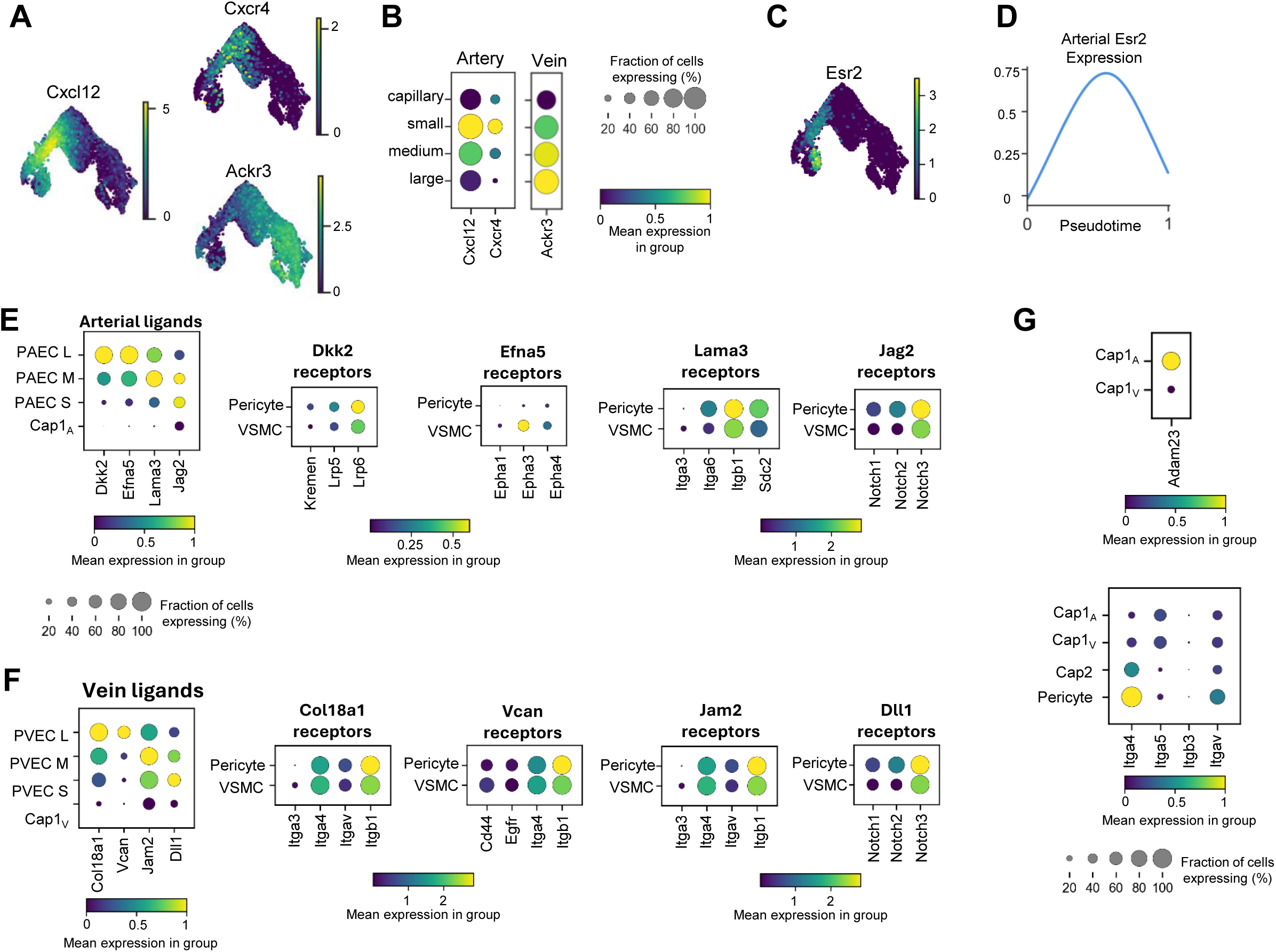
Transcriptomic determination of vessel size provides biologic insight into endothelial derived cellular communication. (A) UMAPs depicted gene expression (low to high: purple to yellow) of *Cxcl12* and its two receptors, *Cxcr4* and *Ackr3*. (B) Dotplots showing *Cxcl12* and *Cxcr4* in arteries by vessel size, and *Ackr3* in veins by vessel size. Relative gene expression is indicated by color (low to high, purple to yellow). Proportion of population expressing the gene is indicated by the size of the circle. (C) UMAPs colored by gene expression of *Esr2* (low to high: purple to yellow). (D) Histogram of *Esr2* gene expression across the arterial pseudotime branch. (E) Dotplots of select ligands expressed by PAEC exhibiting differential expression by vessel size, and corresponding dotplots of receptor expression for each ligand in mural cells. Relative gene expression is indicated by color (low to high: purple to yellow). (F) Dotplots of select ligands expressed by PVEC exhibiting differential expression by vessel size, and corresponding dotplots of receptor expression for each ligand in mural cells. Relative gene expression is indicated by color (low to high: purple to yellow). Proportion of population expressing the gene is indicated by the size of the circle for (E) and (F). (G) Dotplots of *Adam23* expression in Cap1_A_ and Cap1_V_, and *Adam23* receptor expression in cells within vascular cell types in the alveolar niche. Relative gene expression is indicated by color (low to high: purple to yellow). Proportion of population expressing the gene is indicated by the size of the circle

Thus, we examined whether the spatial distribution of the estrogen receptors, *Esr1* and *Esr2* aligned with *Cxc12*-*Cxcr4* enrichment in small arteries. *Esr1* showed limited expression in vascular EC **(SFig. 3A-C)**. In contrast, *Esr2* was restricted to the PAEC, with highest expression in EC from small and medium arteries (**Fig. 4C**), peaking in arterial EC with intermediate pseudotime values, similar to the expression pattern of *Cxcl12* and *Cxcr4* (**Fig. 4D**).

We next investigated whether size based transcriptomic assignment of EC could identify putative cell-cell communication enriched at specific locations along the pulmonary vascular tree. We identified ligands that were differentially expressed in small versus large PAEC and determined the expression of putative receptors for each ligand in mural cells (**Fig. 4E**). Large PAEC highly expressed *Dkk2*, and *Efna5*, a high affinity ligand for *Epha3*, highly expressed in a subset of VSMC. PAEC from medium sized arteries highly express *Lama3*, a component of the basement membrane protein, laminin 5, likely promoting cell adhesion to mural cells highly expressing known interacting partners such as *Itgb1* and *Sdn2*^46,47^. Arterial EC from small and medium arteries selectively express the Notch ligand *Jag2*, a pathway shown to drive angiogenesis particularly in response to hypoxia^48^. Using a similar approach, we also identified location-specific differences in cell-cell communication originating from PVEC (**Fig. 4F**). EC from large veins highly expressed *Col18a1*, the gene encoding the anti-angiogenic protein, endostatin, and *Vcan* encoding versican^49^. Both ligands can bind to integrins expressed by mural cells, and Versican-CD44 interactions likely support mural cell adhesion^50^.The Ig-superfamily adhesion receptor, *Jam2*, was highly expressed by PVEC from medium sized veins, and small PVEC highly express the Notch ligand *Dll1*, potentially signaling to *Notch3*-expressing pericytes to promote vascular stabilization^51^.

Spatially resolved cellular communication even revealed unique putative cellular communication along the arterial and venous sides of the capillaries **(Fig. 4G)**. For example, the atypical ADAM, *Adam23*, which promotes cell adhesion, is only expressed by the arterial Cap1 EC^33^. Taken together, these data highlight that assigning vessel sizes across endothelial cells can identify physiologically relevant and distinct ligand-receptor interactions at specific segments of the pulmonary circulation.

### Vessel-size analytical framework reveals size-dependent responses to vascular injury

We next evaluated whether this analytical framework could provide insights into the pulmonary vascular response to injuries that may not be uniform across the pulmonary circulation. Exposure of neonatal mice to chronic hyperoxia disrupts pulmonary vascular growth and pathologic vascular remodeling. We and others have previously reported that in contrast to suppressing proliferation of Cap1 EC, hyperoxia induces a proliferative response in PVEC^3,52^. However, it was unknown if this proliferative response was predominant in specific portions of the venous circulation. Examining the expression of a panel of cell-cycle genes confirmed heightened expression of pro-proliferation genes on the venous side of the pulmonary circulation, with peak expression in Cap1_V_ (**Fig. 5A).** By calculating proliferation scores among PVEC from different sized vessels, we found that the majority of normoxic EC exhibited low proliferation scores, with a small group of cells from each sized vein demonstrating higher scores. In response to hyperoxia, a greater number of PVEC from small and medium veins exhibited high proliferation scores (**Fig. 5B**). This was associated with a significant increase in top proliferation associated genes in venous-Cap1 and PVEC from small and medium veins (**Fig. 5C**). We validated these findings, using *in vivo* EdU incorporation assays **(Figure 5D)**. Smaller veins had the greatest percent proliferation, and the percent of EdU^+^ PVEC negatively correlated with increasing diameter (**Fig. 5E**). Specifically, hyperoxia only significantly increased proliferation in PVEC in veins smaller than 50μm in diameter (**Fig. 5F**).

**Figure 5:**
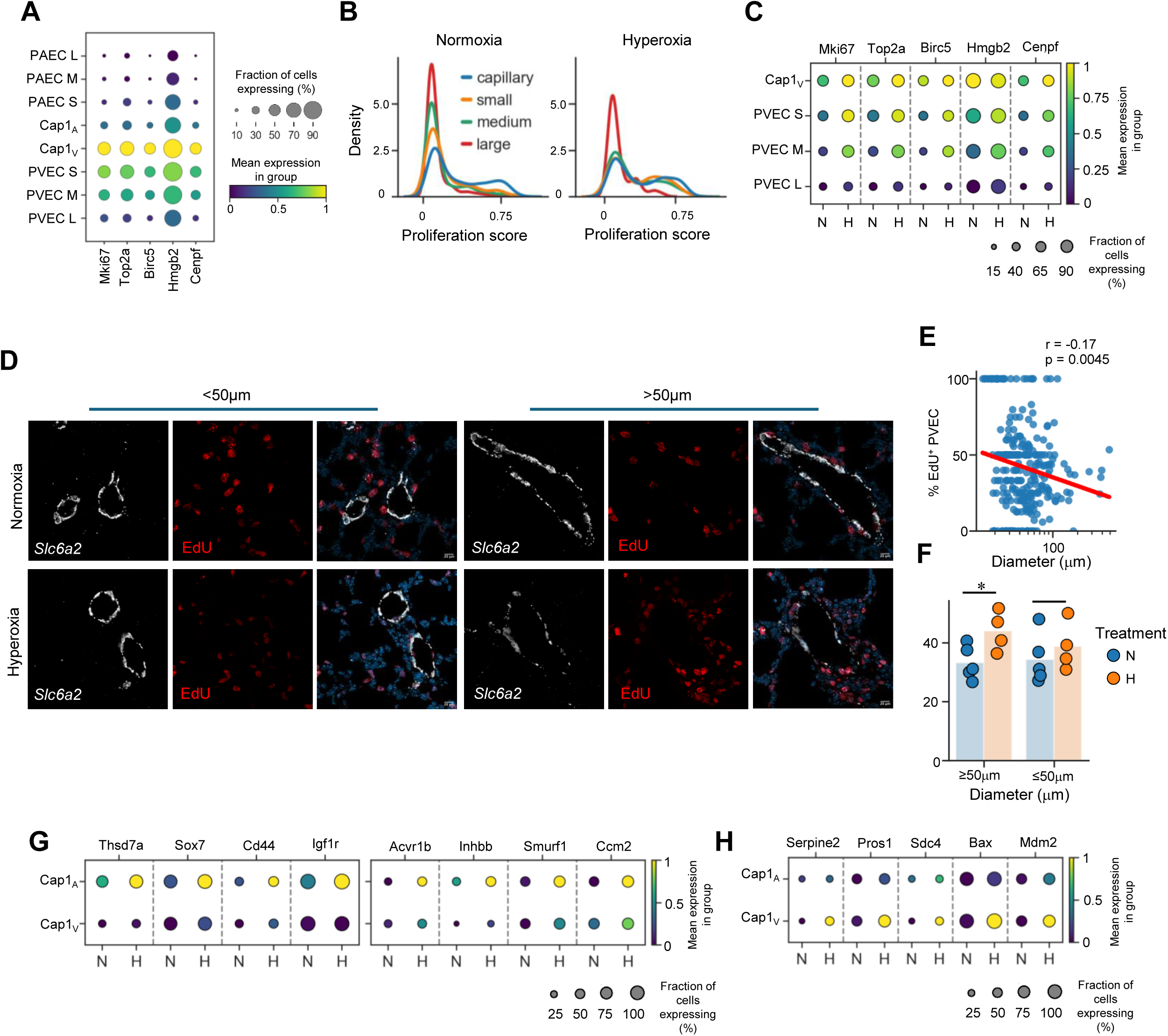
Vessel-size analytical framework reveals size-dependent responses to vascular injury. (A) Dotplot of select proliferation marker genes in EC by vessel type and size. Relative gene expression is indicated by color (low to high: purple to yellow), and proportion of population expressing each gene is indicated by the size of the circle. (B) Kernel density estimate showing distribution of proliferation scores of PVEC by vessel size in normoxia and hyperoxia. (C) Dotplot of select proliferation marker genes in PVEC by vessel size in normoxia and hyperoxia. Relative gene expression is indicated by color (low to high: white to red), and proportion of population expressing each gene is indicated circle size. (D) Representative image of *in situ* hybridization to detect expression of the venous marker gene *Slc6a2* (white) and EdU (Red) in lung tissue from mice at P3 in either normoxia or hyperoxia. Calibration bar=. (E) Quantification of the proportion of EdU+ PVECs by vessel size. Each point represents a vessel. A total of 265 vessels were imaged across normoxic and hyperoxic mice (n=4-5). Line in red is a linear regression on the log10 normalized vessel size, r and p value are from Pearson’s correlation. (F) Quantification of EdU+ PVECs binned by vessel size, less than or equal to or greater than 50 μm. Each point represents a mouse where PVEC nuclei and EdU+ PVEC nuclei counts were summed for each vessel size bin and total %EdU+ PVEC was calculated. The bar represents the mean of the mice from each condition. *p=0.036 by student’s t-test.

We next examined whether hyperoxia induced distinct transcriptomic alterations of Cap1 EC located on either the arterial or venous side of the circulation. In the Cap1_A_ EC, hyperoxia increased genes associated with angiogenesis and TGFβ signaling (**Fig. 5G**). For example, hyperoxia increased *Thsd7a* a secreted glycoprotein that promotes filopodia formation and EC migration, *Sox7*, a Sox family member essential for vasculogenesis and hypoxia-induced angiogenesis, *Cd44*, a hyaluronan receptor that promotes EC proliferation, and *Igf1r*, which promotes stabilization of nascent blood vessels^53–56^. Hyperoxia also increased a number of genes that regulate BMP/TGFβ signaling including up-regulation of *Acvr1b* and *Inhbb* in the Cap1_A_ EC, which encode Alk4 and activin B, a pathway implicated in pathologic pulmonary vascular remodeling, and *Smurf1*, an additional molecule that shifts homeostatic BMP signaling toward TGFβ-mediated remodeling^57,58^. Hyperoxia induced a distinct gene signature in Cap1_V_ EC (**Fig. 5H**), increasing the expression of *Serpine2* and *Pros1,* regulators of coagulation that have also been shown to regulate vascular remodeling and barrier function^59,60^. Hyperoxia also increased *Bax*, a pro-apoptotic regulator induced by stress, and *Mdm2*, a pro-survival factor that inhibits p53 signaling, a known pathway driving hyperoxia induced lung injury. Taken together these data suggest that Cap1 are comprised of two distinct subsets marked by gene signatures delineating their arterial versus venous localization and exhibiting divergent responses to hyperoxia-driven vascular injury.

### Transcriptomic gradients to identify endothelial cells by vessel size are conserved across development and species

To determine the generalizability and external validity of our vessel size framework, we applied it to five additional lung-specific single-cell or nucleus RNAseq datasets spanning multiple ages, disease states, and species. These included two murine datasets: one overlapping with our developmental timepoints and experimental conditions^61^ (**Fig. 6A**) and one from adult lung^62^ (**Fig. 6B**); and three human datasets: a neonatal lung dataset from the first day of life^63^ (**Fig. 6C**), a BPD cohort from 1 month to 3 years of age^64^ (**Fig. 6D**), and an adult lung dataset^65^ (**Fig. 6E**). We used the same approach as above, re-embedding PAEC, Cap1, and PVEC and utilizing pseudotime to approximate vessel size. We found that the terminal ends of the venous and arterial clusters exhibited the highest size scores, which gradually declined toward central Cap1 cluster enabling segmentation of vessels by approximate size (**Supplemental Fig. 4A-D**). Notably, we found that many of the size-related genes were consistent including *Eln,* which was highest in the largest vessels, and *Col4a1,* highest in the capillaries. Integration of these datasets and the present data demonstrate that cell type labels are conserved and embed together, even with multiple species and age cohorts **(Fig. 6E)**. Despite computing vessel size scores on each individual dataset, this score gradient is conserved in the integrated embedding. Using the developmental mouse dataset most similar to our own we see that many of the expression patterns, including size specific-gene expression, cell signaling, and changes in hyperoxia are also observable in this dataset^61^ **(Supplemental Fig. 5 A-D)**.

**Figure 6:**
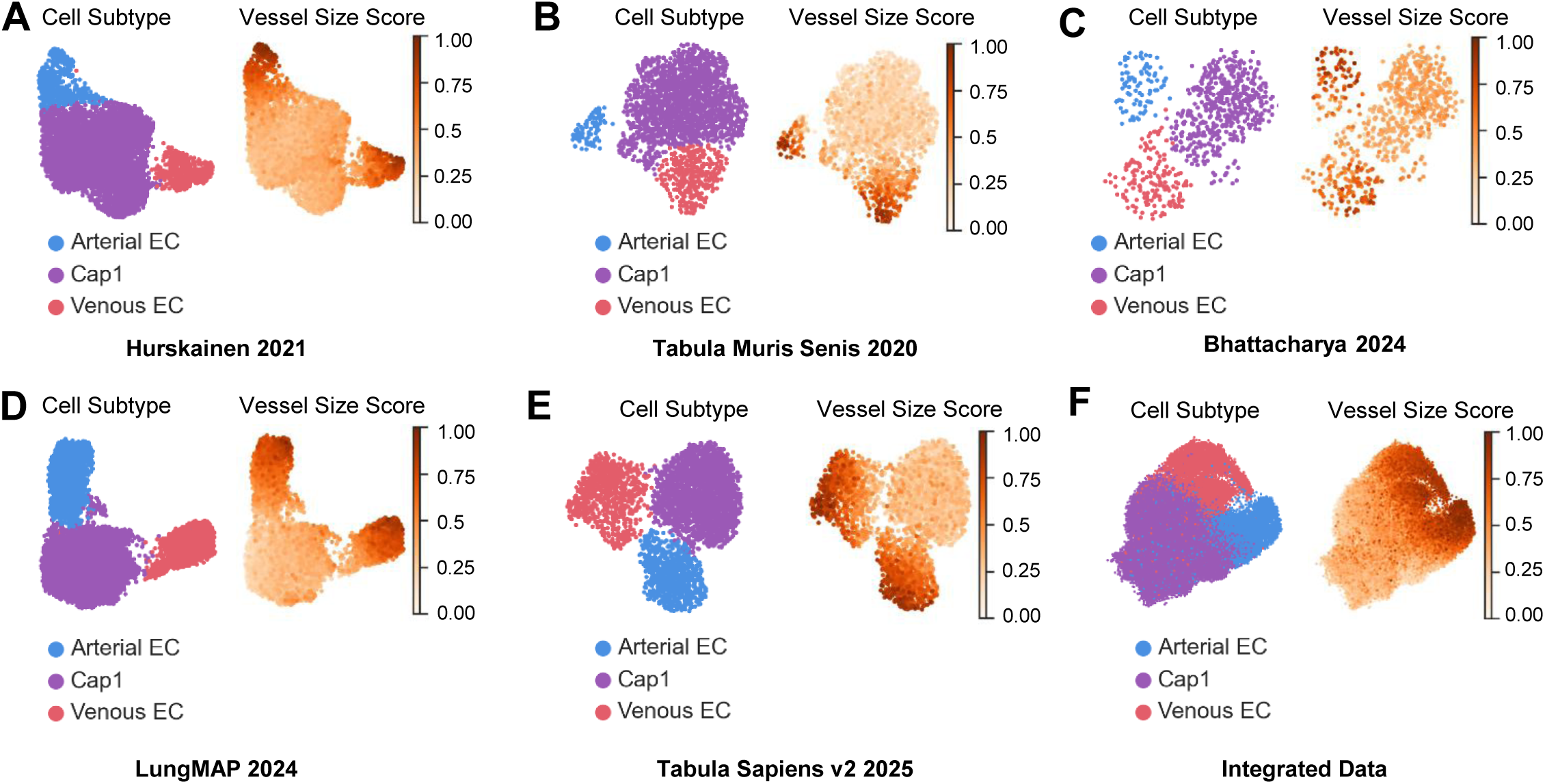
Transcriptomic gradients to identify endothelial cells by vessel size are conserved across development and species. UMAPs of EC cell subtype and vessel scores from datasets derived from (A) the developing mouse lung exposed to normoxia or hyperoxia^61^; (B) the adult mouse lung^62^; (C) one day old infant^63^ (D) control and BPD infants^64^; and (E) adult human lung^65^. UMAP of integrated datasets’ vascular endothelial cells vessel size scoring (low to high, white to orange).

Together, these data demonstrate the reproducibility and consistency of this framework for attributing vessel-size scores to individual arterial, venous, and Cap1 EC across heterogenous sequencing datasets. This reproducibility is maintained despite variations in tissue preparation, sample sources, library preparation, and sequencing technology, highlighting that transcriptional continuums expressed along the vascular hierarchy are conserved across lifespan and species.

## Discussion

Single cell transcriptomics has revealed robust EC heterogeneity but linking these molecular profiles to vascular architecture remains a major challenge. By relating gene expression to vessel size scores, we uncovered evolutionarily and developmentally conserved transcriptional gradients corresponding to vessel size and polarization along the arterial-venous axis. This method offers a scalable and biologically meaningful approach to stratify EC across the pulmonary vascular structural continuum. ^61–65^ Our framework also demonstrates conserved transcriptomic gradients across the pulmonary vascular tree across datasets, ages, disease states, and species, demonstrating highly generalizable and broadly reproducible vessel size-associated transcriptional patterns.

Although all pulmonary EC share core transcriptional programs reflective of their common lineage, their identities are further shaped by cues from the local microenvironment (e.g. oxygen tension, flow dynamics, paracrine signals from adjacent cells). Our data shows that these differing cues produce a transcriptional continuum of gene expression rather than gene expression signatures restricted to separate endothelial subtypes. The main physiologic functions of discrete portions of the pulmonary circulation change across the vascular bed: pressure regulation in arteries, gas exchange in capillaries, and reservoir capacity in veins. By aligning gene expression patterns to inferred vessel size, our framework enables the identification of mechanisms directing regionally-distinct endothelial functions within the lung. For example, *Foxo1*, a FOXO family transcription factor required for vascular growth and patterning in embryonic development was highly expressed by arterial and venous EC of medium sized vessels, highlighting its role in promoting quiescence of maturing vessels in this context^66,67^. The heightened expression of muscarinic receptors in small and medium sized veins likely promotes NO-mediated vasodilation, maintaining the obligatory low vascular tone on the post-capillary side of the circulation required to provide the transpulmonary gradient required for adequate pulmonary blood flow^68^. Examination of endothelial subtypes at greater resolution also allows the identification of gene expression by specialized subsets that would be obscured by ensemble averaging. For example, our data show that a small subset of Cap1 express the neurotrophin receptor *Ntrk2* during normal development, a gene previously reported to be only induced in Cap1 EC after injury^34,69^. Whether these *Ntrk2*+ arterial Cap1 represents a specialized subset with the capacity that preferentially expands in response to injury remains to be determined.

Many pulmonary vascular diseases induce endothelial dysfunction and vascular remodeling in select segments of the pulmonary circulation. For example, PAH is characterized by preferential muscularization of small arterioles, BPD often involves microvascular rarefication followed by aberrant angiogenesis, and pulmonary veno-occlusive disease (PVOD) preferentially affects small veins and venules^13^. To identify molecular mechanisms most relevant to specific vascular pathologies, focusing on the transcriptional and cell signaling aberrations occurring at the level of the vasculature most affected has the greatest chance of identifying disease specific, novel therapies. For example, although it is common for groups to report cell-cell communication across all cells within a single cell dataset, it is more likely that biologically relevant cellular communication occurs within specialized multi-cellular niches that are distinct at different levels of the circulation. This was clear in our dataset, which localized canonical pathways regulating vascular development and stability to discrete portions of the circulation, including highest expression of *Efna5* in large PAEC, potentially modulating the contractility of surrounding VSMC to regulate vascular tone, and *Jag2* and *Dll1* expression by small PAEC and PVEC, respectively, likely promoting pericyte differentiation to promote vascular patterning and stabilization of nascent vessels^70–72^.

We also demonstrated that our analytical framework allowed the identification of differential responses of pulmonary EC derived from different sized vessels to injury. Reports from our group and others have shown VEC proliferate in response to diverse lung injuries and we accurately localized the venous proliferative response to small veins^3,52^. We also found that by delineating the arterial and venous polarization of Cap1, we could identify enrichment of unique biologic responses in each subgroup in response to hyperoxia. In the Cap1_A_, hyperoxia up-regulated genes that regulate angiogenesis and drive TGFβ-mediated vascular remodeling, where on the opposite side of the alveolus, the Cap1_V_ exhibited a heightened injury response with up-regulation of genes modulating coagulation, and p53-mediated signaling, likely reflecting a response to the markedly increased paO_2_ experienced by these cells.

Our study has several limitations. First, we made somewhat arbitrary divisions of cells in relation to vessel size, there is no ground-truth within the transcriptomic data relative to the diameter of a ‘small’ or ‘large’ vessel. Our analytic framework also performed optimally on datasets containing a high sampling of ECs with high sequencing depth across the vascular hierarchy, with less cells and/or lower sequencing depth offering less resolution. Finally, we limited this analysis to establishing the transcriptional gradients expressed along the endothelial hierarchy. However, VSMC and pericytes also exist along the same vascular continuum and it is likely that the mural cells would exhibit similar transcriptomic gradients. Identification of mural cell location from the transcriptome would further refine cell-cell communication predictions, particularly when considering either ligands or receptors that are membrane bound.

Together these results highlight the power of applying a vessel-sized score to the interpretation of transcriptomic data. Signaling gradients are fundamental to organogenesis, providing spatial cues that guide cell fate decisions and tissue organization. Thus, it is unsurprising that ECs exhibit a transcriptomic continuum reflecting their position along physiologic axes. By relating gene expression to physiologically relevant locations, this approach revealed organized zones of endothelial specialization with potential clinical and therapeutic relevance. Although spatial transcriptomics and other *in situ* methods offer high-resolution views of gene expression within tissue, they are costly and low-throughout, restricted to predefined regions or gene panels, and still limited in their ability to resolve transcriptional gradients at cellular resolution across a complex hierarchy such as the vasculature. In contrast, our framework allows inference of spatial positioning directly from single cell transcriptomic data, enabling thousands of cells to be aligned along an anatomic continuum. In the future, additional physiological parameters such as blood oxygen concentration, flow dynamics, or stretch, could be integrated as complementary axes to further refine interpretations of transcriptomic changes in the pulmonary endothelium. Further, incorporating multi-modal datasets, such as single cell ATAC-seq, could illuminate the regulatory mechanisms underlying these gradients to refine our understanding of EC fate and phenotype transitions. We also anticipate that this strategy could be extended to other tissues and cells that exist along similar continua defined by size or other relevant gradients, with a similar paradigm already suggested in the brain^73^. Finally, applying this strategy to diverse datasets may enable resolution of common hierarchical regulatory systems driving key molecular mechanisms directing cell-fate, organ development, and disease.

## Acknowledgments

None

## Author contributions

S.N.S. contributed to study design, data collection, data analysis, interpretation, and writing of the manuscript.

C.K. contributed to study design, data analysis, interpretation, and writing of the manuscript.

F.Z. contributed to data analysis, interpretation, and writing of the manuscript.

D.N.C. contributed to data interpretation and writing of the manuscript.

C.M.A. supervised all aspects of the project, including the design, data analysis and interpretation, contributed to the writing and final approval of the manuscript.

---

**Supplemental Figure 1:**
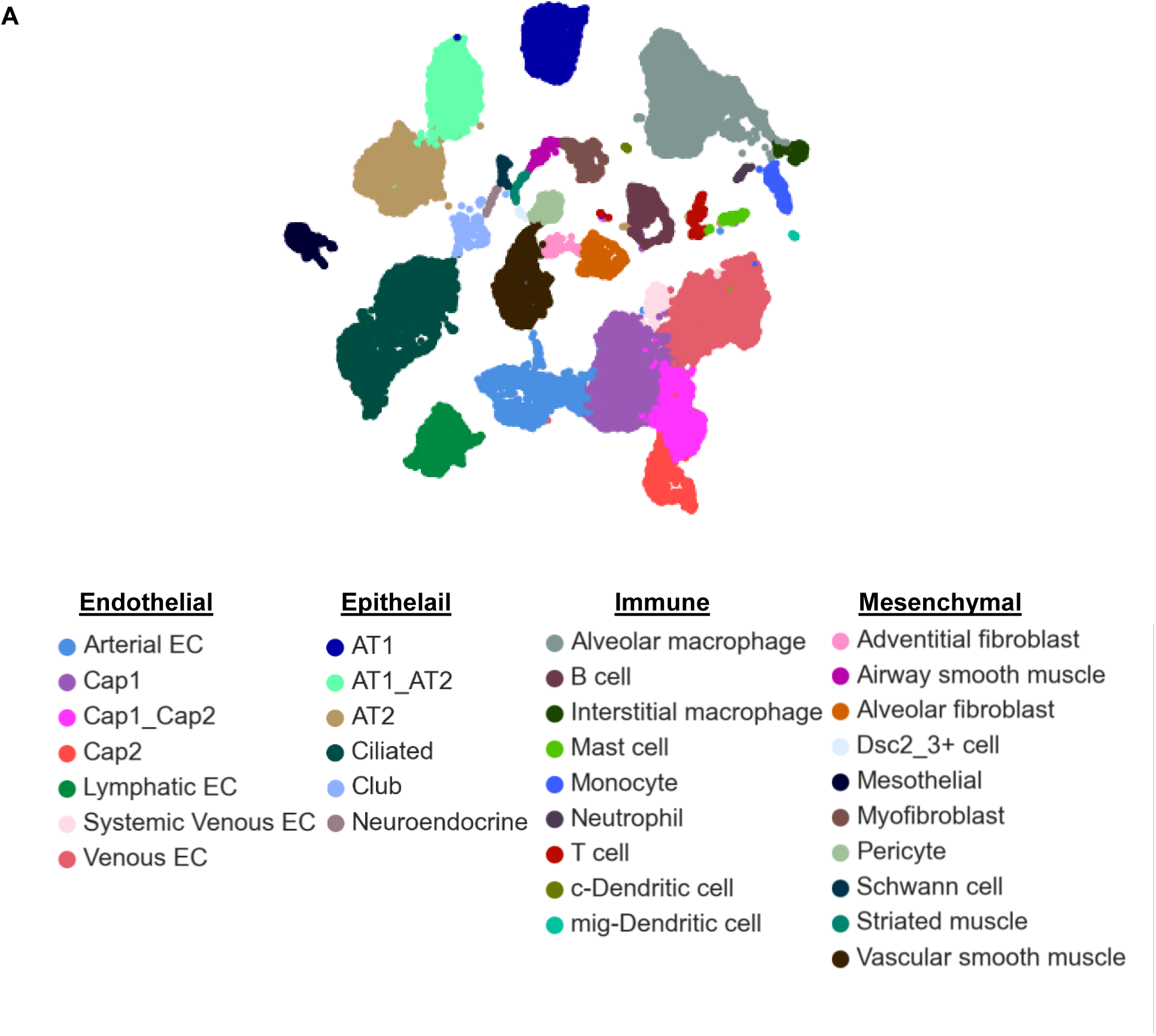
Cell types detected in P3 murine dataset. (A) UMAP of P3 mouse dataset colored by cell type

**Supplemental Figure 2:**
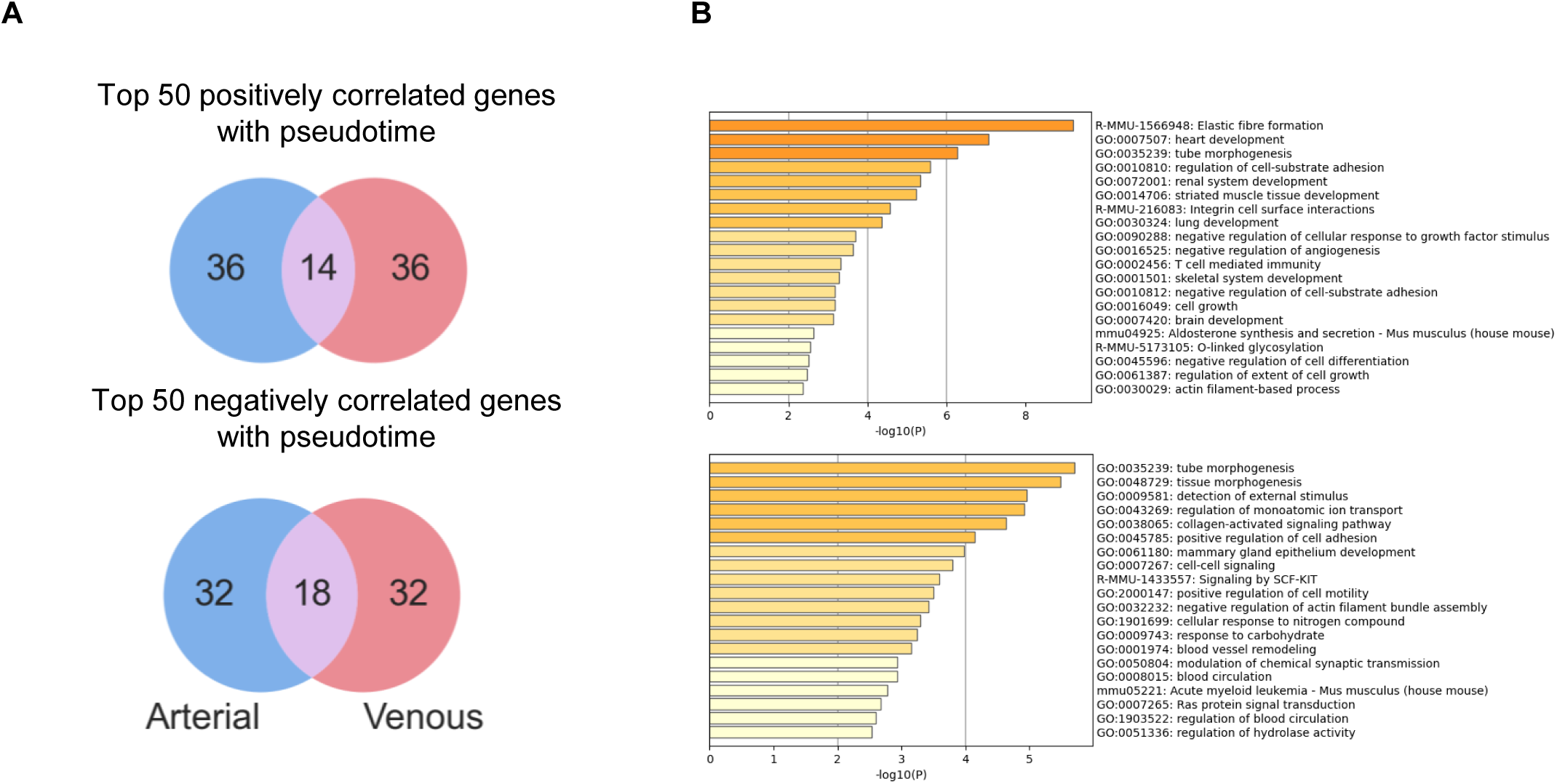
Shared genes and pathways correlated with pseudotime in arteries and veins. (A) Venn diagrams showing overlap of top 50 genes correlated with pseudotime in Arterial and Venous EC branches. (B) All pathways enriched in shared genes between artery and vein negatively (top) and positively (bottom) correlated with pseudotime

**Supplemental Figure 3:**
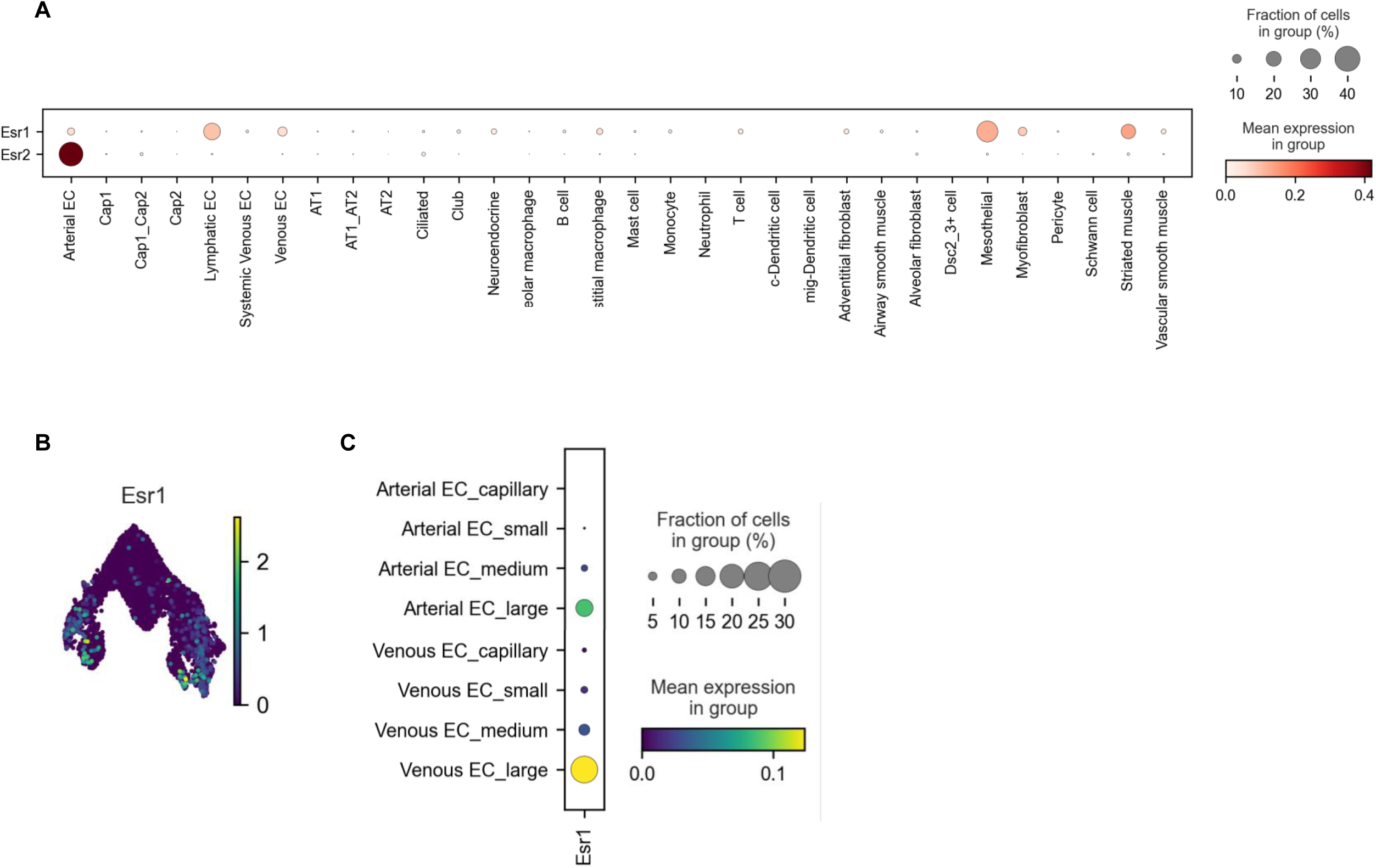
Estrogen receptor expression in P3 murine dataset. (A) Dotplot of Esr1 and Esr2 expression across all cell types in P3 murine dataset. Average gene expression is indicated by color, in log counts per ten thousand, (low to high, white to red). Proportion of population expressing the gene is indicated by the size of the circle. (B) UMAP of vascular endothelial cells colored by Esr1 expression (low to high, purple to yellow). (C) Dotplot showing gene expression with distinct localization along the vascular hierarchy in PAECs and PVECs. Average gene expression is indicated by color (low to high, purple to yellow). Proportion of population expressing the gene is indicated by the size of the circle.

**Supplemental Figure 4:**
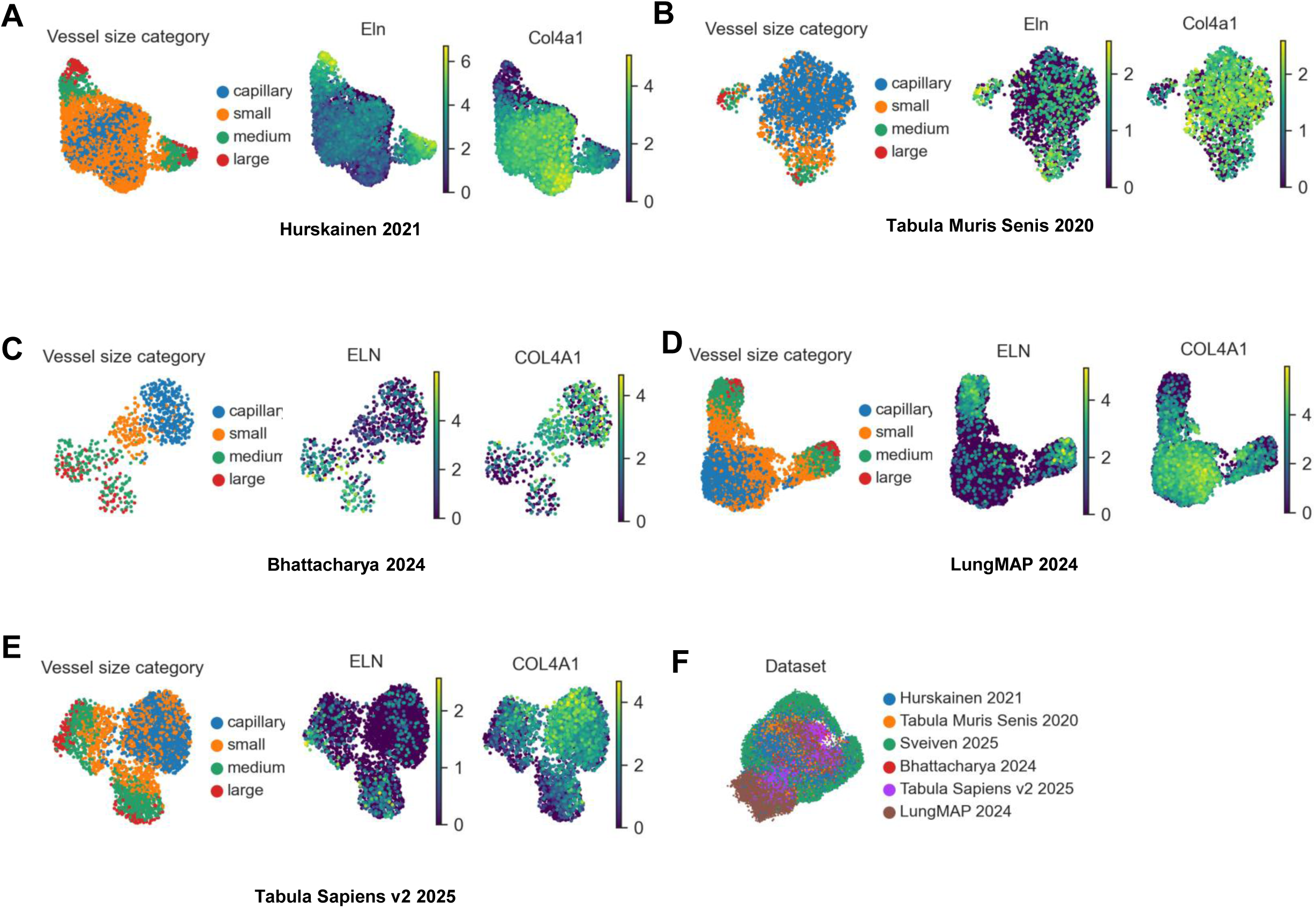
Vessel size scoring of mouse and human vascular endothelial cells across development. (A) UMAP of neonatal mouse vascular endothelial cells colored by cell type, vessel size category and gene gene expression (low to high, purple to yellow). of *Eln*, and *Col4a1*. (B) UMAP of adult mouse vascular endothelial cells colored by cell type, vessel size category and gene gene expression (low to high, purple to yellow). of *Eln*, and *Col4a1*. (C) UMAP of neonatal human vascular endothelial cells colored by cell type, vessel size category and gene gene expression (low to high, purple to yellow). of *ELN*, and *COL4A1*. (D) UMAP of 1-month-3 year old human vascular endothelial cells colored by cell type, vessel size category and gene gene expression (low to high, purple to yellow). of *ELN*, and *COL4A1*. (E) UMAP of adult human vascular endothelial cells colored by cell type, vessel size category and gene gene expression (low to high, purple to yellow). of *ELN*, and *COL4A1*. (F) UMAP of integrated data colored by dataset

**Supplemental Figure 5:**
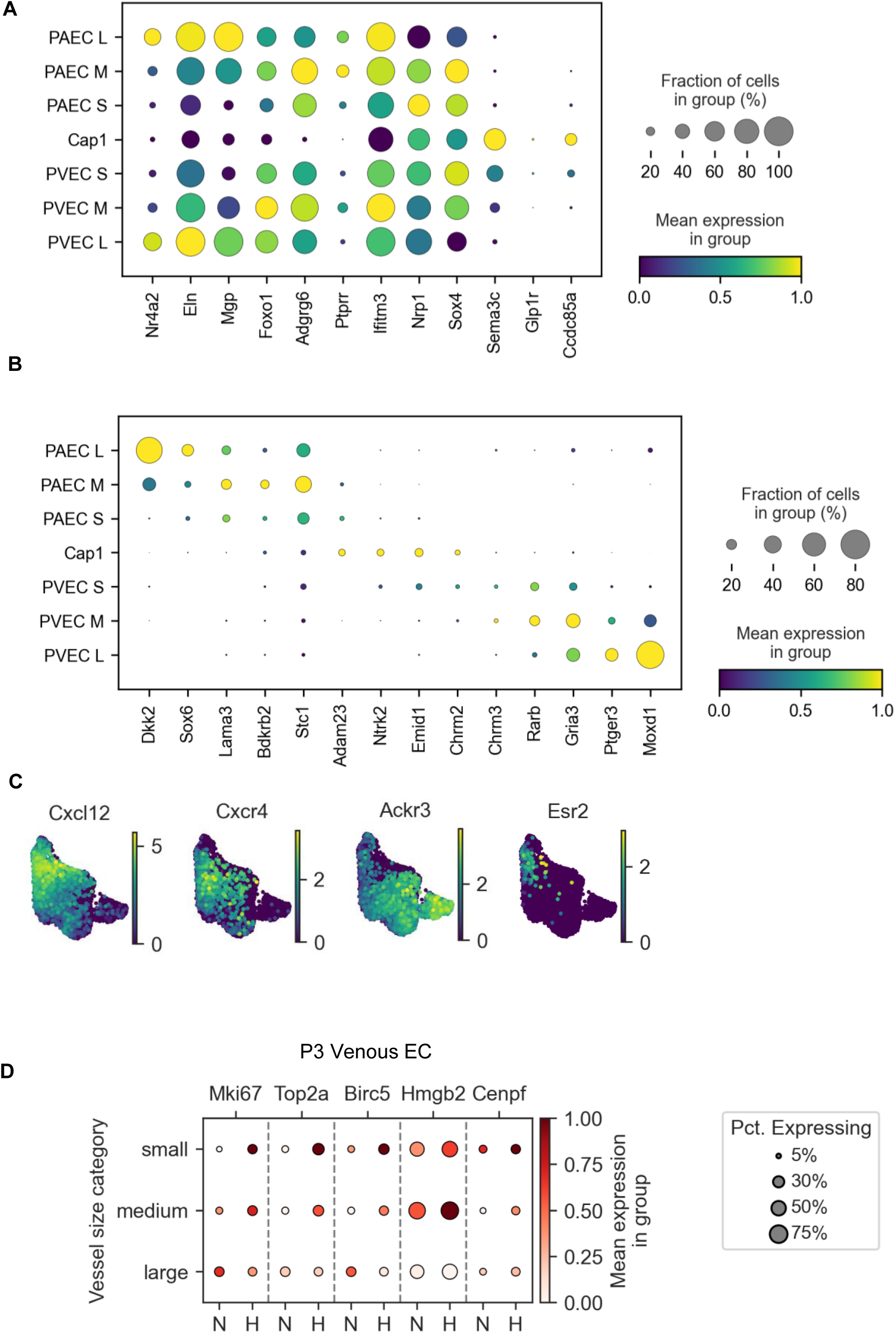
Vessel size transcriptomic patterns shown in Hurskainen et al scRNA-seq dataset. (A) Dotplot showing gene expression associated with size along the vascular hierarchy in Hurskainen et al.. Average gene expression is indicated by color (low to high, purple to yellow). Proportion of population expressing the gene is indicated by the size of the circle. (B) Dotplot showing gene expression with distinct localization along the vascular hierarchy in Hurskainen et al Average gene expression is indicated by color (low to high, purple to yellow). Proportion of population expressing the gene is indicated by the size of the circle. (C) UMAP of vascular endothelial cells of signaling genes *Cxcl12*, *Cxcr4*, *Ackr3*, and *Esr2* from Hurskainen et al colored by gene expression (low to high, purple to yellow). (D) Dotplot of select proliferation marker genes in P3 PVECs by vessel size in normoxia and hyperoxia. Average gene expression is indicated by color (low to high, white to red). Proportion of population expressing the gene is indicated by the size of the circle.

## References

1. Gillich A, Zhang F, Farmer CG, Travaglini KJ, Tan SY, Gu M, Zhou B, Feinstein JA, Krasnow MA, Metzger RJ. Capillary cell-type specialization in the alveolus. Nature. 2020;586:785–789.

2. Schupp JC, Adams TS, Cosme C, Raredon MSB, Yuan Y, Omote N, Poli S, Chioccioli M, Rose K-A, Manning EP, et al. Integrated Single-Cell Atlas of Endothelial Cells of the Human Lung. Circulation. 2021;144:286–302.

3. Zanini F, Che X, Knutsen C, Liu M, Suresh NE, Domingo-Gonzalez R, Dou SH, Zhang D, Pryhuber GS, Jones RC, et al. Developmental diversity and unique sensitivity to injury of lung endothelial subtypes during postnatal growth. iScience. 2023;26:106097.

4. Zepp JA, Morley MP, Loebel C, Kremp MM, Chaudhry FN, Basil MC, Leach JP, Liberti DC, Niethamer TK, Ying Y, et al. The genomic, epigenomic, and biophysical cues controlling the emergence of the lung alveolus. Science. 2021;371:eabc3172.

5. Zepp JA, Morrisey EE. Cellular crosstalk in the development and regeneration of the respiratory system. Nat Rev Mol Cell Biol. 2019;20:551–566.

6. Vila Ellis L, Cain MP, Hutchison V, Flodby P, Crandall ED, Borok Z, Zhou B, Ostrin EJ, Wythe JD, Chen J. Epithelial Vegfa Specifies a Distinct Endothelial Population in the Mouse Lung. Dev Cell. 2020;52:617–630.e6.

7. Heumos L, Schaar AC, Lance C, Litinetskaya A, Drost F, Zappia L, Lücken MD, Strobl DC, Henao J, Curion F, et al. Best practices for single-cell analysis across modalities. Nat Rev Genet. 2023;24:550–572.

8. Sharma A, Niethamer TK. Specialized Pulmonary Vascular Cells in Development and Disease. Annual Review of Physiology. 2025;87:229–255.

9. Ardini-Poleske ME, Clark RF, Ansong C, Carson JP, Corley RA, Deutsch GH, Hagood JS, Kaminski N, Mariani TJ, Potter SS, et al. LungMAP: The Molecular Atlas of Lung Development Program. Am J Physiol Lung Cell Mol Physiol. 2017;313:L733–L740.

10. Kalucka J, de Rooij LPMH, Goveia J, Rohlenova K, Dumas SJ, Meta E, Conchinha NV, Taverna F, Teuwen L-A, Veys K, et al. Single-Cell Transcriptome Atlas of Murine Endothelial Cells. Cell. 2020;180:764–779.e20.

11. Stevens T, Garcia JGN, Shasby DM, Bhattacharya J, Malik AB. Mechanisms regulating endothelial cell barrier function. American Journal of Physiology-Lung Cellular and Molecular Physiology [Internet]. 2000 [cited 2025 Dec 22];Available from: https://journals.physiology.org/doi/10.1152/ajplung.2000.279.3.L419

12. Hansmann G, Sallmon H, Roehr CC, Kourembanas S, Austin ED, Koestenberger M. Pulmonary hypertension in bronchopulmonary dysplasia. Pediatr Res. 2021;89:446–455.

13. Montani D, Lau EM, Dorfmüller P, Girerd B, Jaïs X, Savale L, Perros F, Nossent E, Garcia G, Parent F, et al. Pulmonary veno-occlusive disease. European Respiratory Journal. 2016;47:1518–1534.

14. Young MD, Behjati S. SoupX removes ambient RNA contamination from droplet-based single-cell RNA sequencing data. GigaScience. 2020;9:giaa151.

15. Wolf FA, Angerer P, Theis FJ. SCANPY: large-scale single-cell gene expression data analysis. Genome Biology. 2018;19:15.

16. Wolock SL, Lopez R, Klein AM. Scrublet: Computational Identification of Cell Doublets in Single-Cell Transcriptomic Data. Cell Syst. 2019;8:281–291.e9.

17. Characterization of cell fate probabilities in single-cell data with Palantir | Nature Biotechnology [Internet]. [cited 2025 May 9];Available from: https://www.nature.com/articles/s41587-019-0068-4

18. La Manno G, Soldatov R, Zeisel A, Braun E, Hochgerner H, Petukhov V, Lidschreiber K, Kastriti ME, Lönnerberg P, Furlan A, et al. RNA velocity of single cells. Nature. 2018;560:494–498.

19. Bergen V, Soldatov RA, Kharchenko PV, Theis FJ. RNA velocity—current challenges and future perspectives. Molecular Systems Biology. 2021;17:e10282.

20. Stirling DR, Swain-Bowden MJ, Lucas AM, Carpenter AE, Cimini BA, Goodman A. CellProfiler 4: improvements in speed, utility and usability. BMC Bioinformatics. 2021;22:433.

21. Alvira CM. Aberrant Pulmonary Vascular Growth and Remodeling in Bronchopulmonary Dysplasia. Frontiers in Medicine [Internet]. 2016 [cited 2022 Nov 4];3. Available from: https://www.frontiersin.org/articles/10.3389/fmed.2016.00021

22. Somekawa S, Imagawa K, Hayashi H, Sakabe M, Ioka T, Sato GE, Inada K, Iwamoto T, Mori T, Uemura S, et al. Tmem100, an ALK1 receptor signaling-dependent gene essential for arterial endothelium differentiation and vascular morphogenesis. Proceedings of the National Academy of Sciences. 2012;109:12064–12069.

23. Brengle BM, Lin M, Roth RA, Jones KD, Wagenseil JE, Mecham RP, Halabi CM. A new mouse model of elastin haploinsufficiency highlights the importance of elastin to vascular development and blood pressure regulation. Matrix Biol. 2023;117:1–14.

24. Luo G, Ducy P, McKee MD, Pinero GJ, Loyer E, Behringer RR, Karsenty G. Spontaneous calcification of arteries and cartilage in mice lacking matrix GLA protein. Nature. 1997;386:78–81.

25. Van Tiel CM, De Vries CJM. NR4All in the vessel wall. The Journal of Steroid Biochemistry and Molecular Biology. 2012;130:186–193.

26. Musa G, Cazorla-Vázquez S, van Amerongen MJ, Stemmler MP, Eckstein M, Hartmann A, Braun T, Brabletz T, Engel FB. Gpr126 (Adgrg6) is expressed in cell types known to be exposed to mechanical stimuli. Annals of the New York Academy of Sciences. 2019;1456:96–108.

27. Wilhelm K, Happel K, Eelen G, Schoors S, Oellerich MF, Lim R, Zimmermann B, Aspalter IM, Franco CA, Boettger T, et al. FOXO1 couples metabolic activity and growth state in the vascular endothelium. Nature. 2016;529:216–220.

28. Carra S, Foglia E, Cermenati S, Bresciani E, Giampietro C, Lora Lamia C, Dejana E, Beltrame M, Cotelli F. Ve-ptp Modulates Vascular Integrity by Promoting Adherens Junction Maturation. PLoS ONE. 2012;7:e51245.

29. Sun X, Zeng H, Kumar A, Belser JA, Maines TR, Tumpey TM. Constitutively Expressed IFITM3 Protein in Human Endothelial Cells Poses an Early Infection Block to Human Influenza Viruses. J Virol. 2016;90:11157–11167.

30. Maden CH, Fantin A, Ruhrberg C. Neuropilin ligands in vascular and neuronal patterning. Biochem Soc Trans. 2009;37:1228–1232.

31. Hallmann R, Horn N, Selg M, Wendler O, Pausch F, Sorokin LM. Expression and Function of Laminins in the Embryonic and Mature Vasculature. Physiological Reviews. 2005;85:979–1000.

32. Fernandes L, Fortes ZB, Nigro D, Tostes RCA, Santos RAS, Catelli de Carvalho MH. Potentiation of Bradykinin by Angiotensin-(1-7) on Arterioles of Spontaneously Hypertensive Rats Studied In Vivo. Hypertension. 2001;37:703–709.

33. Cal S, Freije JMP, López JM, Takada Y, López-Otín C. ADAM 23/MDC3, a Human Disintegrin That Promotes Cell Adhesion via Interaction with the αvβ3 Integrin through an RGD-independent Mechanism. MBoC. 2000;11:1457–1469.

34. Kong CSL, V M, Pantaleón-García J, Evans SE, Chen J. Truncated NTRK2 is induced in CAP1 endothelial cells during mouse lung injury-repair. iScience. 2025;28:112973.

35. Kawata T, Muramatsu K, Shishito N, Ichikawa-Tomikawa N, Oishi T, Kakuda Y, Akiyama Y, Yamaguchi K, Sakamoto M, Sugino T. EMID1, a multifunctional molecule identified in a murine model for the invasion independent metastasis pathway. Sci Rep. 2021;11:16372.

36. Sailem HZ, Al Haj Zen A. Morphological landscape of endothelial cell networks reveals a functional role of glutamate receptors in angiogenesis. Sci Rep. 2020;10:13829.

37. Saternos HC, Forero KV, Meqdad MA, Buqaileh R, Sunderman CL, Gallagher G, Messer WS, Mohieldin AM, Mucci CA, Kumariya S, et al. Muscarinic acetylcholine receptor 3 localized to primary endothelial cilia regulates blood pressure and cognition. Sci Rep. 2025;15:3745.

38. Cho SJ, George CLS, Snyder JM, Acarregui MJ. Retinoic acid and erythropoietin maintain alveolar development in mice treated with an angiogenesis inhibitor. Am J Respir Cell Mol Biol. 2005;33:622–628.

39. Yun EJ, Lorizio W, Seedorf G, Abman SH, Vu TH. VEGF and endothelium-derived retinoic acid regulate lung vascular and alveolar development. American Journal of Physiology-Lung Cellular and Molecular Physiology. 2016;310:L287–L298.

40. Chandrasekaran P, Negretti NM, Sivakumar A, Liberti DC, Wen H, Peers de Nieuwburgh M, Wang JY, Michki NS, Chaudhry FN, Kaur S, et al. CXCL12 defines lung endothelial heterogeneity and promotes distal vascular growth. Development. 2022;149:dev200909.

41. Yi D, Liu B, Wang T, Liao Q, Zhu MM, Zhao Y-Y, Dai Z. Endothelial Autocrine Signaling through CXCL12/CXCR4/FoxM1 Axis Contributes to Severe Pulmonary Arterial Hypertension. Int J Mol Sci. 2021;22:3182.

42. Klouda T, Tsikis ST, Hirsch TI, Kim Y, Li Y, Gaal J, Zhao Z, Friehs I, Shyy JY-J, Raby BA, et al. Smooth muscle Cxcl12 contributions to vascular remodeling in flow and hypoxia-induced pulmonary hypertension. J Biol Chem. 2025;301:110207.

43. Bordenave J, Thuillet R, Tu L, Phan C, Cumont A, Marsol C, Huertas A, Savale L, Hibert M, Galzi J-L, et al. Neutralization of CXCL12 attenuates established pulmonary hypertension in rats. Cardiovasc Res. 2020;116:686–697.

44. Sauvé K, Lepage J, Sanchez M, Heveker N, Tremblay A. Positive feedback activation of estrogen receptors by the CXCL12-CXCR4 pathway. Cancer Res. 2009;69:5793–5800.

45. Chen X, Austin ED, Talati M, Fessel JP, Farber-Eger EH, Brittain EL, Hemnes AR, Loyd JE, West J. Estrogen Inhibition Reverses Pulmonary Arterial Hypertension and Associated Metabolic Defects. Eur Respir J. 2017;50:1602337.

46. Utani A, Nomizu M, Matsuura H, Kato K, Kobayashi T, Takeda U, Aota S, Nielsen PK, Shinkai H. A unique sequence of the laminin alpha 3 G domain binds to heparin and promotes cell adhesion through syndecan-2 and-4. J Biol Chem. 2001;276:28779–28788.

47. Ryan MC, Lee K, Miyashita Y, Carter WG. Targeted disruption of the LAMA3 gene in mice reveals abnormalities in survival and late stage differentiation of epithelial cells. J Cell Biol. 1999;145:1309–1323.

48. Pietras A, von Stedingk K, Lindgren D, Påhlman S, Axelson H. JAG2 induction in hypoxic tumor cells alters Notch signaling and enhances endothelial cell tube formation. Mol Cancer Res. 2011;9:626–636.

49. O’Reilly MS, Boehm T, Shing Y, Fukai N, Vasios G, Lane WS, Flynn E, Birkhead JR, Olsen BR, Folkman J. Endostatin: an endogenous inhibitor of angiogenesis and tumor growth. Cell. 1997;88:277–285.

50. Wu YJ, La Pierre DP, Wu J, Yee AJ, Yang BB. The interaction of versican with its binding partners. Cell Res. 2005;15:483–494.

51. Tefft JB, Bays JL, Lammers A, Kim S, Eyckmans J, Chen CS. Notch1 and Notch3 coordinate for pericyte-induced stabilization of vasculature. Am J Physiol Cell Physiol. 2022;322:C185–C196.

52. Wong J, Zhao G, Adams-Tzivelekidis S, Wen H, Chandrasekaran P, Michki SN, Gentile ME, Singh M, Kass-Gergi S, Mendoza M, et al. Dynamic behavior and lineage plasticity of the pulmonary venous endothelium. Nat Cardiovasc Res. 2024;3:1584–1600.

53. Kuo M-W, Wang C-H, Wu H-C, Chang S-J, Chuang Y-J. Soluble THSD7A is an N-glycoprotein that promotes endothelial cell migration and tube formation in angiogenesis. PLoS One. 2011;6:e29000.

54. Klomp J, Hyun J, Klomp JE, Pajcini K, Rehman J, Malik AB. Comprehensive transcriptomic profiling reveals SOX7 as an early regulator of angiogenesis in hypoxic human endothelial cells. J Biol Chem. 2020;295:4796–4808.

55. Savani RC, Cao G, Pooler PM, Zaman A, Zhou Z, DeLisser HM. Differential involvement of the hyaluronan (HA) receptors CD44 and receptor for HA-mediated motility in endothelial cell function and angiogenesis. J Biol Chem. 2001;276:36770–36778.

56. Jacobo SMP, Kazlauskas A. Insulin-like growth factor 1 (IGF-1) stabilizes nascent blood vessels. J Biol Chem. 2015;290:6349–6360.

57. Guignabert C, Savale L, Boucly A, Thuillet R, Tu L, Ottaviani M, Rhodes CJ, De Groote P, Prévot G, Bergot E, et al. Serum and Pulmonary Expression Profiles of the Activin Signaling System in Pulmonary Arterial Hypertension. Circulation. 2023;147:1809–1822.

58. Zhu H, Kavsak P, Abdollah S, Wrana JL, Thomsen GH. A SMAD ubiquitin ligase targets the BMP pathway and affects embryonic pattern formation. Nature. 1999;400:687–693.

59. Madjene C, Boutigny A, Bouton M-C, Arocas V, Richard B. Protease Nexin-1 in the Cardiovascular System: Wherefore Art Thou? Front Cardiovasc Med. 2021;8:652852.

60. Joussaume A, Kanthou C, Pardo OE, Karayan-Tapon L, Benzakour O, Dkhissi F. The Vitamin K-Dependent Anticoagulant Factor, Protein S, Regulates Vascular Permeability. Curr Issues Mol Biol. 2024;46:3278–3293.

61. Hurskainen M, Mižíková I, Cook DP, Andersson N, Cyr-Depauw C, Lesage F, Helle E, Renesme L, Jankov RP, Heikinheimo M, et al. Single cell transcriptomic analysis of murine lung development on hyperoxia-induced damage. Nat Commun. 2021;12:1565.

62. Almanzar N, Antony J, Baghel AS, Bakerman I, Bansal I, Barres BA, Beachy PA, Berdnik D, Bilen B, Brownfield D, et al. A single-cell transcriptomic atlas characterizes ageing tissues in the mouse. Nature. 2020;583:590–595.

63. Bhattacharya S, Myers JA, Baker C, Guo M, Danopoulos S, Myers JR, Bandyopadhyay G, Romas ST, Huyck HL, Misra RS, et al. Single-Cell Transcriptomic Profiling Identifies Molecular Phenotypes of Newborn Human Lung Cells. Genes (Basel). 2024;15:298.

64. LungMAP - Dataset [Internet]. [cited 2025 Jul 9];Available from: https://www.lungmap.net/dataset/?experiment_id=LMEX0000004400

65. Quake SR, Consortium TTS. Tabula Sapiens reveals transcription factor expression, senescence effects, and sex-specific features in cell types from 28 human organs and tissues [Internet]. 2024 [cited 2025 Apr 20];2024.12.03.626516. Available from: https://www.biorxiv.org/content/10.1101/2024.12.03.626516v1

66. Furuyama T, Kitayama K, Shimoda Y, Ogawa M, Sone K, Yoshida-Araki K, Hisatsune H, Nishikawa S, Nakayama K, Nakayama K, et al. Abnormal angiogenesis in Foxo1 (Fkhr)-deficient mice. J Biol Chem. 2004;279:34741–34749.

67. Andrade J, Shi C, Costa ASH, Choi J, Kim J, Doddaballapur A, Sugino T, Ong YT, Castro M, Zimmermann B, et al. Control of endothelial quiescence by FOXO-regulated metabolites. Nat Cell Biol. 2021;23:413–423.

68. Furchgott RF, Zawadzki JV. The obligatory role of endothelial cells in the relaxation of arterial smooth muscle by acetylcholine. Nature. 1980;288:373–376.

69. Niethamer TK, Planer JD, Morley MP, Babu A, Zhao G, Basil MC, Cantu E, Frank DB, Diamond JM, Nottingham AN, et al. Longitudinal single-cell profiles of lung regeneration after viral infection reveal persistent injury-associated cell states. Cell Stem Cell [Internet]. 2025 [cited 2025 Jan 17];0. Available from: https://www.cell.com/cell-stem-cell/abstract/S1934-5909(24)00441-7

70. Zhang Z, Tremblay J, Raelson J, Sofer T, Du L, Fang Q, Argos M, Marois-Blanchet F-C, Wang Y, Yan L, et al. EPHA4 regulates vascular smooth muscle cell contractility and is a sex-specific hypertension risk gene in individuals with type 2 diabetes. J Hypertens. 2019;37:775–789.

71. Nadeem T, Bogue W, Bigit B, Cuervo H. Deficiency of Notch signaling in pericytes results in arteriovenous malformations. JCI Insight. 2020;5:e125940, 125940.

72. Wang Y, Pan L, Moens CB, Appel B. Notch3 establishes brain vascular integrity by regulating pericyte number. Development. 2014;141:307–317.

73. Vanlandewijck M, He L, Mäe MA, Andrae J, Ando K, Del Gaudio F, Nahar K, Lebouvier T, Laviña B, Gouveia L, et al. A molecular atlas of cell types and zonation in the brain vasculature. Nature. 2018;554:475–480.

